# Multimodal Protein Retrieval via Joint Representation Learning from Sequences and Cryo-EM Density Maps

**DOI:** 10.64898/2026.09.02.749033

**Authors:** Akshat Tulsani, Aditya Maddur Guruprakash, Skanda Shreesha Prasad

## Abstract

Aligning protein sequences with cryo-EM density maps remains challenging due to limited paired data, structural heterogeneity, varying map resolutions, and the presence of multiple conformational states. In this work, we propose a multimodal representation learning framework that learns a shared latent space between protein sequences and cryo-EM density maps for cross-modal retrieval.

Our approach combines pretrained protein sequence embeddings with a volumetric cryo-EM encoder trained using self-supervised representation learning and transfer learning. The resulting model enables bidirectional retrieval between sequences and density maps while learning biologically meaningful structural representations.

Experimental results demonstrate strong retrieval performance across both sequence-to-map and map-to-sequence tasks, achieving median retrieval ranks of 2–3 within a database of 3,275 cryo-EM maps. The learned embedding space shows a clear separation between matched and unmatched sequence–map pairs and remains robust across varying cryo-EM resolutions. Additionally, the model generalizes across species, successfully retrieving conserved mouse protein structures using human sequence embeddings. Our findings demonstrate that joint latent-space learning provides a promising direction for connecting protein sequences with cryo-EM structural representations, with potential applications in structural retrieval, protein annotation, and multimodal biological representation learning.

## 1. Introduction

Understanding the relationship between protein sequence and three-dimensional structure is a fundamental challenge in molecular biology. Protein language models such as *ESM-2* learn powerful representations from amino-acid sequences, capturing evolutionary and biochemical information [22, 25]. Meanwhile, cryo-electron microscopy (cryo-EM) provides experimentally derived threedimensional density maps that reveal the geometric structure of biomolecules [7, 17, 23]. Despite the availability of both modalities, most machine learning methods treat sequence and structural information independently.

Sequence-based models operate purely on amino-acid sequences, while cryo-EM pipelines focus on reconstructing atomic models from density maps. Similarly, structure prediction frameworks such as *AlphaFold* [16], *RoseTTAFold* [2], and *ESMFold* [22] infer three-dimensional structures from sequences without directly incorporating experimental cryo-EM observations during representation learning. Although resources such as AlphaFold DB [30], UniProt [27], UniRef [26], and EMDataBank [19] provide large-scale sequence and structure-related data, these information sources are typically exploited separately.

As a result, existing approaches learn complementary but disconnected representations of proteins. A unified multimodal representation learning framework that jointly models protein sequences and cryo-EM structural images could bridge this gap. Such a framework could enable models to learn shared latent representations capturing the relationship between sequence composition and three-dimensional structural geometry.

These multimodal representations could support several downstream tasks including sequence-to-structure retrieval, protein identification from cryo-EM maps, mutation impact analysis, and structural clustering. More broadly, they could improve our ability to understand how protein sequences give rise to functional molecular structures.

### Motivation: Cross-Modal Protein Retrieval

Currently, identifying a protein in a raw cryo-EM density map often requires manual “atomic model building,” a process that can take days or weeks. Ideally, a scientist should be able to query a cryo-EM map and instantly retrieve candidate protein sequences whose structures match the observed density. Achieving this requires learning a shared representation that links protein sequences with experimental structural observations.

In this work, we propose a multimodal framework that learns joint representations of protein sequences and cryo-EM density maps through predictive latent objectives inspired by Joint Embedding Predictive Architectures (JEPA) [1, 20].

## 2. Related Work

### ESM-2 (2023)

Traditional bioinformatics methods rely on computationally expensive multiple sequence alignments (MSAs) to infer protein properties. To bypass this bottleneck, a pure Transformer architecture is used to scale to 15 billion parameters, trained unsupervised on hundreds of millions of evolutionary sequences. This large-scale pre-training allows the model to implicitly capture the biophysical rules of protein folding, enabling accurate zero-shot prediction of 3D structural contacts directly from its internal self-attention maps. [22].

### ModelAngelo (2023)

In the domain of experimental structural biology, interpreting cryo-electron microscopy (cryo-EM) density maps historically required manual tracing of atomic coordinates. To automate this reconstruction process, this work proposes a hybrid architecture combining Graph Neural Networks (GNNs) and Transformers. By formulating structure determination as a graph classification task and conditioning the network on amino acid sequences, the methodology successfully extracts local structural motifs to generate high-resolution atomic models. [15].

### CLIP (2021)

To bridge disparate data modalities into a unified representational space, this work introduces a massive-scale dual-encoder architecture originally designed for vision-language alignment. The framework employs an InfoNCE contrastive loss to maximize the cosine similarity of paired embeddings while minimizing it for negative samples within a batch. This approach established the standard paradigm for zero-shot cross-modal retrieval. However, contrastive objectives are notoriously sensitive to data quality and require massive batches of perfectly aligned high-fidelity pairs. [24].

### I-JEPA (2023)

The inherent instability of contrastive learning on noisy datasets necessitates robust alternatives that do not rely on negative sampling or extensive data augmentation. This work introduces a non-contrastive framework that predicts the latent representation of a masked target region conditioned on an unmasked context. By enforcing learning through latent prediction rather than pairwise contrast, the architecture avoids representation collapse and demonstrates high computational efficiency. Although originally evaluated on 2D images, the framework’s resilience to corrupted or missing input features makes it an ideal methodology for handling the weak pairing and sparsity inherent in experimental cryo-EM datasets [1].

### CryoDRGN (2021)

While traditional cryo-EM pipelines reconstruct a single static 3D density map, many biological macromolecules exhibit continuous structural flexibility. To capture this heterogeneity, this work introduces a deep generative model based on Variational Autoencoders (VAEs) to learn a continuous latent representation of cryo-EM particle images. By mapping 2D projection images into a shared low-dimensional latent space, the architecture successfully reconstructs a continuous trajectory of 3D density maps, proving that neural networks can extract meaningful latent manifolds from highly noisy electron microscopy data. [34].

### Foldseek (2023)

As the number of predicted and experimental 3D structures exploded, the bioinformatics community required scalable retrieval systems to search massive structural databases. To bypass the extreme computational cost of 3D spatial alignment, this work discretizes physical 3D geometries into a 1D structural alphabet, allowing structural search to be executed using ultra-fast sequence alignment heuristics. This methodology drastically reduced structure-to-structure retrieval times, establishing the standard baseline for large-scale structural database querying. Despite its speed, the algorithm strictly operates on clean, high-resolution atomic coordinates (such as those in the PDB or AlphaFold DB); it cannot process or retrieve unmodeled, voxel-based raw density maps, limiting its utility on noisy experimental cryo-EM data [29].

### OneProt (2025)

The exponential growth of structural databases has motivated the development of unified retrieval systems across biological modalities. This work proposes a multimodal protein foundation model utilizing a parameter-efficient, adapter-based fine-tuning strategy to align primary sequences, 3D structures, and textual annotations into a shared latent space. The framework establishes current state-of-the-art benchmarks for cross-modal protein retrieval (e.g., Recall@k, MRR), validating the efficacy of latent search mechanisms for biological discovery. Nevertheless, the model relies on structural embeddings derived from clean, high-resolution atomic coordinates rather than raw experimental density maps, highlighting a critical gap that our proposed voxel-grounded predictive architecture aims to resolve [10].

## 3. Methodology

We introduce a multimodal representation learning framework for aligning protein sequences and cryo-EM density maps within a shared latent space. Our approach combines pretrained protein language models, cryo-EM-specific volumetric representation learning, and cross-modal predictive objectives to enable bidirectional retrieval between sequence and structure representations.

An overview of the proposed framework is shown in Figure 1. The pipeline consists of three major stages: (1) selfsupervised pretraining of the cryo-EM volume encoder using Cryo2Struct density maps, (2) cross-modal JEPA training on mouse sequence–structure pairs, and (3) transfer learning to the human EMDB dataset for downstream retrieval.

**Figure 1.**
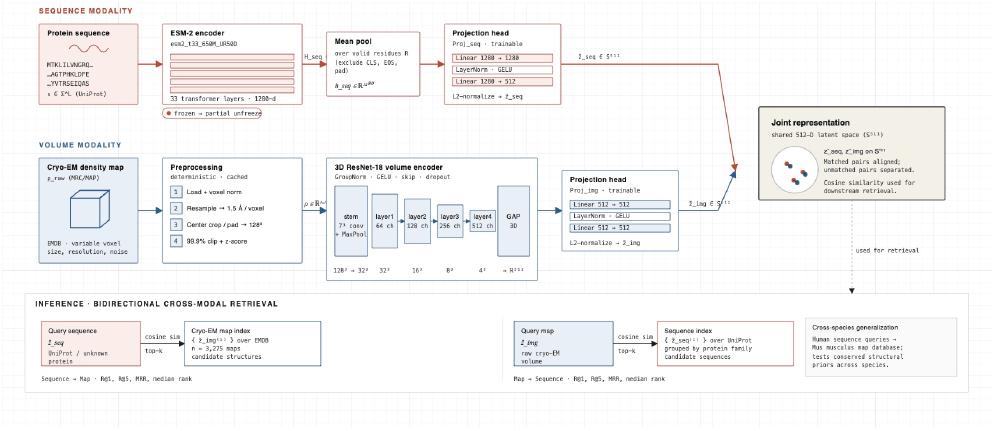
Overview of the proposed multimodal cryo-EM representation learning framework. Cryo2Struct volumes are first used for MAE pretraining of the cryo-EM volume encoder. The model is then trained using cross-modal JEPA objectives on mouse sequence–structure pairs before transfer learning to the human EMDB dataset for bidirectional sequence–map retrieval.

A natural formulation for multimodal sequence– structure learning would be to use a CLIP-style contrastive framework that directly aligns paired protein sequences and cryo-EM maps [24]. However, cryo-EM datasets present several challenges that make strict instance-level contrastive learning difficult.

First, experimentally paired sequence–structure datasets remain relatively limited. While billions of protein sequences are available, only a small subset have experimentally resolved structures and even fewer have associated cryo-EM density maps [19, 27]. Second, cryo-EM data contains substantial structural ambiguity: a single protein sequence may exist across multiple conformational states, while a single cryo-EM map may contain heterogeneous assemblies, multiple proteins, or partially resolved structures [9, 34]. Finally, cryo-EM maps vary significantly in reconstruction quality, resolution, and noise characteristics, making direct one-to-one contrastive alignment unreliable [7, 23].

To address these limitations, we adopt a Joint Embedding Predictive Architecture (JEPA)-style framework that learns predictive relationships between modalities through latent-space prediction objectives rather than relying purely on exact pairwise matching [1, 20]. Our approach is further motivated by broader predictive pretraining paradigms in representation learning and vision-language modeling [31].

Rather than enforcing exact embedding equality between sequence and structure representations, the model learns a shared latent space where protein sequences and cryo-EM maps contain mutually predictive structural information. This allows the framework to better tolerate conformational variability, noisy supervision, and heterogeneous cryo-EM structures while still enabling strong bidirectional retrieval performance.

### 3.1. Data Exploration and Setup

Our primary objective is to learn shared multimodal representations between protein sequences and experimentally derived cryo-EM density maps for *Homo sapiens*. The long-term motivation is to support experimental structural biology pipelines operating in de novo settings, where researchers aim to identify or characterize proteins directly from cryo-EM observations without requiring fully reconstructed atomic models.

In particular, we are interested in enabling cross-modal retrieval between protein sequences and cryo-EM density maps:

- retrieving candidate protein sequences given a cryo-EM density map,
- retrieving structurally compatible cryo-EM maps given a protein sequence,
- and learning latent structural representations that capture sequence–structure relationships across experimental conformations.

To construct the dataset, we curated a subset of entries from the EM Data Bank (EMDB) corresponding to *Homo sapiens*. Each EMDB entry was linked to associated protein sequences through UniProt cross-references and meta-data annotations. The resulting dataset contains over 3,280 cryo-EM density maps together with associated protein sequences information.

#### Dataset Attributes

Each cryo-EM entry consists of:

- A volumetric cryo-EM density map

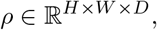

representing experimentally reconstructed electron density
- Resolution metadata describing reconstruction quality (in Å)
- Sample annotations describing biological composition (protein, complex, ligand-bound structure, etc.)
- One or more associated UniProt protein sequences

Unlike conventional image–text datasets used in multi-modal learning, cryo-EM datasets exhibit substantial biological and structural ambiguity. A single cryo-EM map may contain multiple interacting protein subunits, partial assemblies, or heterogeneous conformational states. Similarly, the same protein sequence may appear across multiple cryo-EM maps corresponding to different structural configurations.

These observations highlight several challenges inherent to the dataset:

- **Weak supervision:** Many entries contain incomplete or indirect mappings between sequences and structures.
- **Multi-instance ambiguity:** A single density map may correspond to multiple protein sequences or complexes.
- **Noise and heterogeneity:** Variability in resolution, reconstruction quality, and conformational states reduces the reliability of direct voxel-level comparisons.

Taken together, these properties make strict instance-level alignment unreliable. This motivates the use of predictive latent objectives, where the model learns shared representations across modalities without requiring exact sequence–structure correspondence.

#### 3.1.1. Transfer Learning: House Mouse

In addition to the primary *Homo sapiens* dataset, we also construct a secondary cryo-EM dataset from *Mus musculus* (house mouse) entries in EMDB. The mouse dataset follows the same construction pipeline, linking cryo-EM density maps with associated UniProt protein sequences through EMDB metadata and cross-references. This filtering resulted in a dataset of over 2,517 cryo-EM density maps with associated metadata and protein sequence annotations.

The motivation for including *Mus musculus* is twofold. First, mouse is one of the most widely studied model organisms in structural and molecular biology, resulting in a relatively large number of experimentally resolved cryo-EM structures. Second, mouse and human proteins share strong genetic and structural similarity due to evolutionary conservation [8]. Many core biological pathways, protein families, and structural motifs are conserved across the two species, making mouse-derived structural representations potentially transferable to human protein understanding.

Our experiments therefore explore whether multimodal representations learned from mouse cryo-EM sequence– structure relationships can improve representation learning on the human dataset. Specifically, we investigate a transfer learning setting where the cryo-EM volume encoder and cross-modal alignment objectives are first trained on the *Mus musculus* dataset before being adapted to the *Homo sapiens* dataset.

This setup is particularly relevant because experimentally resolved cryo-EM datasets remain relatively limited compared to sequence-only resources. Cross-species transfer learning may therefore provide a mechanism for improving structural representation learning by leveraging conserved biological structure across organisms.

The transfer learning strategy and training procedure are described further in the Cross-Modal JEPA Training Objective section.

#### 3.1.2. Pretraining: Understanding Cryo-EM Structural Features

Cryo-EM density maps differ substantially from natural images commonly used in representation learning. Unlike RGB images with rich texture, color, and semantic information, cryo-EM maps are single-channel three-dimensional volumetric density fields containing comparatively sparse and low-level structural signals. The structural information is encoded through subtle density variations corresponding to molecular geometry, secondary structure organization, and spatial mass distribution.

This creates an additional challenge for multimodal learning. Before the model can learn meaningful sequence– structure relationships, the volume encoder must first learn to interpret the underlying geometric and structural patterns present in cryo-EM densities. In practice, a randomly initialized 3D encoder spends a significant portion of training learning basic cryo-EM structural primitives such as:

- local density continuity,
- volumetric structural boundaries,
- alpha-helical and beta-sheet patterns,
- spatial organization of macromolecular assemblies,
- and noise characteristics specific to cryo-EM reconstruction pipelines.

To improve structural understanding prior to multimodal alignment, we introduce a dedicated pretraining stage for the cryo-EM volume encoder.

For this stage, we use cryo-EM density maps from the *Cryo2Struct* dataset [11]. The dataset contains approximately 7,900 cryo-EM volumes spanning diverse proteins and structural configurations. These volumes are used exclusively for pretraining the 3D volume encoder to learn generic cryo-EM structural representations before multi-modal sequence alignment.

Importantly, we do not use the primary EMDB multi-modal training dataset during this pretraining phase. The Cryo2Struct dataset is itself derived from a subset of EMDB entries, and using overlapping volumes during both pre-training and multimodal evaluation could introduce leakage between stages. To avoid any possible overlap between pre-training and downstream multimodal retrieval experiments, we keep the Cryo2Struct pretraining volumes fully separate from the EMDB-based sequence–structure alignment dataset.

The goal of this stage is therefore not to learn sequence correspondence, but rather to initialize the volume encoder with stronger structural priors for understanding cryo-EM geometry and density organization. The pretrained encoder is subsequently adapted during cross-modal JEPA training on the human and mouse sequence–structure datasets.

### 3.2. Preprocessing

Cryo-EM density maps obtained from EMDB exhibit significant variability in spatial resolution, voxel scaling, reconstruction quality, and structural extent. To enable stable learning with a 3D neural encoder, we design a preprocessing pipeline that transforms raw cryo-EM density maps into standardized volumetric tensors.

Given a raw cryo-EM map *ρ*_raw_ the preprocessing pipeline consists of four stages:

1. map loading and voxel normalization,
2. spatial resampling,
3. spatial normalization via cropping/padding,
4. intensity normalization.

Cryo-EM, Figure [2], maps are stored as volumetric MRC/MAP files with varying voxel resolutions across experiments. During loading, voxel spacing is extracted directly from the MRC header metadata. In cases where voxel metadata is incomplete or anisotropic, the preprocessing pipeline uses the mean voxel spacing as a stable approximation.

**Figure 2.**
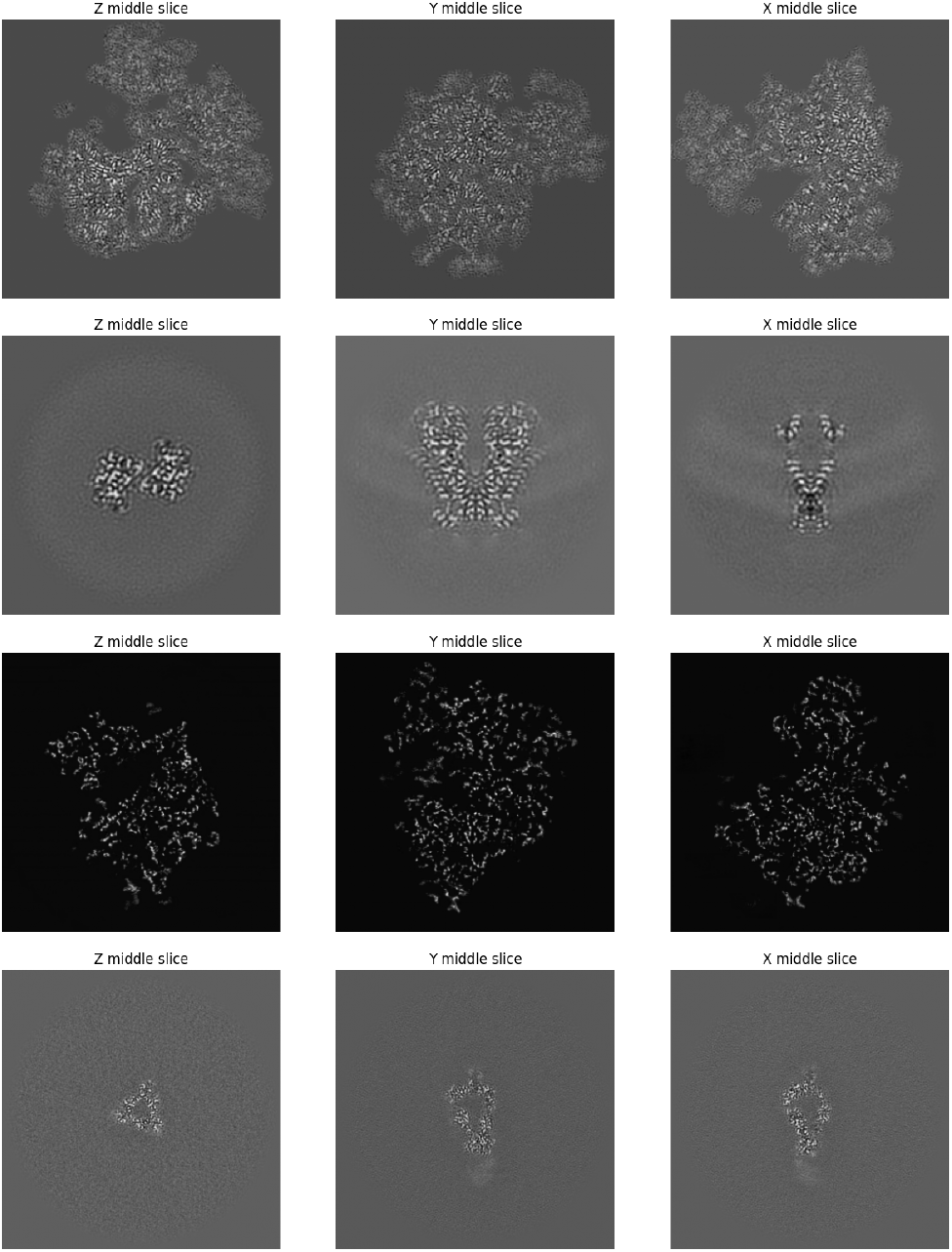
Samples

To ensure geometric consistency across samples, each volume is resampled to a fixed physical resolution of 1.5 Å per voxel using trilinear interpolation:

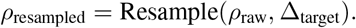

This step ensures that all cryo-EM maps are represented within a consistent spatial coordinate system, allowing the model to learn geometry-aware structural features independent of acquisition settings.

Following resampling, cryo-EM maps vary substantially in spatial dimensions. To obtain fixed-size inputs for the 3D encoder, each volume is center-cropped or zero-padded to a cubic grid:

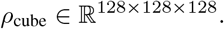

Center cropping and padding preserve the primary structural mass near the center of the volume while enabling uniform tensor dimensions across the dataset.

Cryo-EM, Figure [3], density values vary significantly across experiments due to differences in reconstruction pipelines and noise characteristics. To improve robustness, we apply a two-stage normalization procedure:

- percentile-based clipping (99.9%) to suppress extreme density artefacts,
- z-score normalization computed over non-zero voxels only.

**Figure 3.**
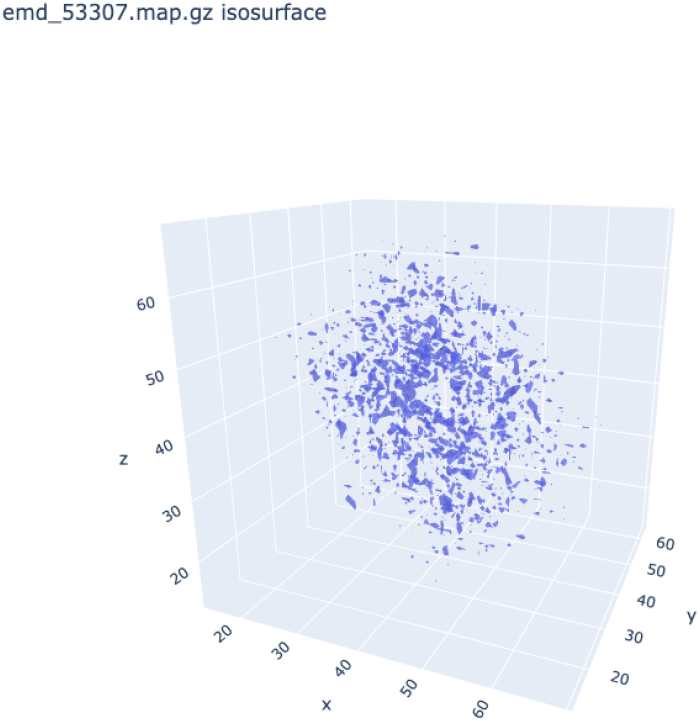
Samples Isosurfaces

Restricting normalization statistics to non-zero voxels prevents zero-padded regions from dominating the density distribution.

The final output of the preprocessing pipeline is a standardized single-channel tensor:

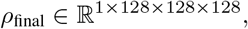

which serves as input to the 3D cryo-EM encoder.

Since volumetric resampling is computationally expensive, all preprocessed maps are cached offline as serialized PyTorch tensors prior to training. This avoids repeated pre-processing during data loading and substantially reduces CPU overhead during multimodal training.

Cryo2Struct pretraining volumes undergo the same spatial normalization and tensor conversion pipeline to maintain consistency with the downstream EMDB multimodal dataset. Since Cryo2Struct maps are already normalized and stored in MRC format, preprocessing primarily consists of voxel resampling, center cropping/padding, and robust z-score normalization before caching.

### 3.3. Encoder

The overall architecuture, Figure 4, consists of two modality-specific encoders that map protein sequences and cryo-EM density maps into a shared latent representation space.

**Figure 4.**
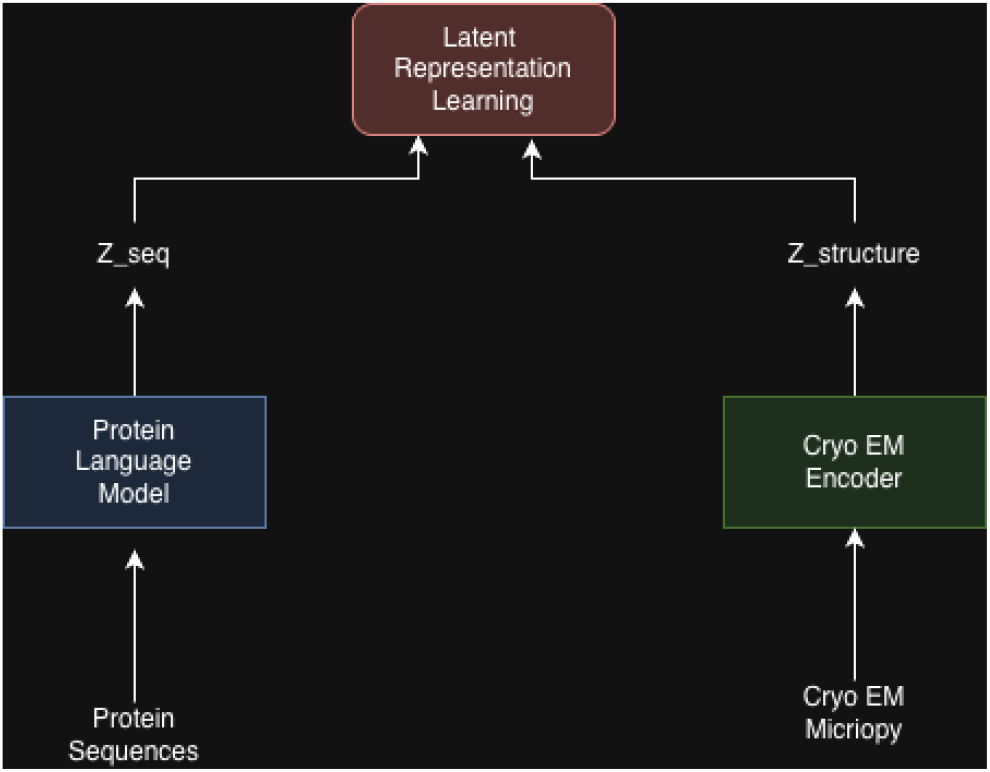
Architecture of the multimodal protein representation model.

#### 3.3.1. Protein Sequence Encoder

Protein sequences are encoded using a pretrained ESM-2 protein language model. In our implementation, we use the `esm2 t33 650M UR50D` checkpoint, which produces 1280-dimensional residue-level representations from the final transformer layer. To preserve pretrained biological and structural knowledge while avoiding overfitting on our relatively small paired dataset, the ESM-2 backbone is kept frozen during the initial training stage. Only a lightweight projection head is trained.

Given a batch of raw amino-acid sequences, the sequence encoder first tokenizes and pads the sequences using the ESM-2 batch converter. The model then extracts final-layer token representations:

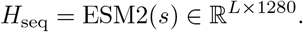

Because ESM-2 is not trained with a dedicated classification-token objective, we do not use the `CLS`token as the sequence summary. Instead, we apply padding-aware mean pooling over real residue positions, excluding `CLS`, `EOS`, and padding tokens:

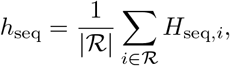

where ℛ denotes valid residue positions.

The pooled sequence embedding is then passed through a two-layer trainable projection head:

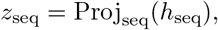

where the projection head maps 1280 *→* 1280 *→* 512 using a linear layer, layer normalization, GELU activation, and a final linear layer.

We retain both the raw projected embedding and its L2-normalized form. The raw projection is used for variance and covariance regularization, while the normalized projection is used for cross-modal alignment and retrieval:

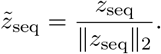

This produces a 512-dimensional sequence representation on the unit hypersphere, matching the latent dimensionality of the cryo-EM volume encoder. During the frozen stage, the ESM-2 forward pass is executed without gradient tracking, reducing memory usage while preserving pre-trained representations. The implementation also supports staged fine-tuning, where the top ESM-2 transformer layers can be unfrozen in later training phases with a smaller learning rate than the projection head.

#### 3.3.2. Volume Encoder

To encode cryo-EM density maps, we employ a 3D convolutional neural network based on a ResNet-18 architecture adapted for volumetric data. The encoder maps a preprocessed density volume

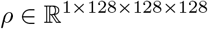

into a compact latent representation aligned with the sequence embedding space.

The volume encoder follows a hierarchical 3D ResNet design. The input volume is first processed by a stem consisting of a 7 *×* 7 *×* 7 convolution with stride 2, followed by Group Normalization and a GELU activation. A max-pooling layer further reduces spatial resolution:

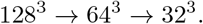

The network then applies four residual stages, each composed of two 3D residual blocks. Spatial resolution is progressively reduced while channel dimensionality increases:

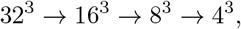

with channel sizes increasing from 64 to 512.

Each residual block consists of two 3 *×* 3 *×* 3 convolutions with skip connections. When spatial resolution or channel dimensions change, the skip path uses a 1 *×* 1 *×* 1 convolution to match dimensions.

We replace Batch Normalization with Group Normalization to improve stability under small batch sizes and heterogeneous input distributions. Cryo-EM volumes vary significantly in resolution and density characteristics, making batch statistics unreliable. Group Normalization decouples normalization from batch composition and yields more stable training.

The final feature map is aggregated using global average pooling:

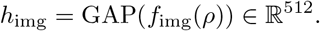

The pooled representation is passed through a two-layer projection head:

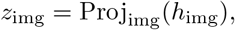

where the projection head maps 512 *→* 512 *→* 512 using a linear layer, layer normalization, GELU activation, and a final linear layer. A dropout layer is applied before projection for regularization.

We retain both the raw projected embedding and its L2-normalized representation. The raw projection is used for variance and covariance regularization objectives, while the normalized embedding is used for cross-modal alignment and retrieval:

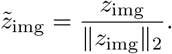

Our design choices are motivated by the characteristics of cryo-EM data:

- **Large receptive fields:** The 7^3^ convolutional stem captures multi-voxel structural patterns such as alpha-helices and beta-sheets.
- **Hierarchical feature extraction:** Progressive downsampling allows the network to capture both local density patterns and global structural organization.
- **Translation robustness:** Global average pooling ensures invariance to small spatial shifts introduced during pre-processing.
- **Regularization:** Dropout and a moderate-capacity backbone help reduce overfitting given the relatively limited number of paired cryo-EM sequence–structure samples. The resulting encoder produces a 512-dimensional embedding on the unit hypersphere aligned with the sequence encoder output, enabling cross-modal predictive learning.

To improve the encoder’s ability to model cryo-EM structural geometry prior to multimodal alignment, we additionally introduce a self-supervised masked autoencoding pretraining stage over Cryo2Struct cryo-EM volumes.

#### 3.3.3. Volume Encoder Pretraining with Cryo-EM MAE

A key challenge in our framework is that the cryo-EM volume encoder must learn meaningful structural features from sparse single-channel 3D density maps. Unlike natural images, cryo-EM maps do not contain color, texture, or object-level semantic cues. Instead, structural information is encoded through local density continuity, secondary-structure-like patterns, global molecular topology, and reconstruction-specific noise. As a result, a randomly initialized 3D encoder may spend much of multimodal training learning basic cryo-EM density features before it can contribute useful gradients to the sequence– structure alignment objective.

To address this, we pretrain the cryo-EM volume encoder using a masked autoencoding objective over Cryo2Struct volumes. The pretraining stage uses approximately 7,390 preprocessed cryo-EM maps from the Cryo2Struct dataset. These volumes are used only for self-supervised volume pretraining and are kept separate from the EMDB sequence–structure alignment dataset to avoid overlap with downstream retrieval evaluation.

The MAE model uses the same 3D ResNet volume backbone as the downstream multimodal model. Given an input volume

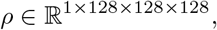

the volume is divided into non-overlapping 3D patches of size 8^3^, producing a 16 *×* 16 *×* 16 patch grid with 4096 total patches. During pretraining, contiguous cubic blocks of patches are masked, and the model is trained to reconstruct the missing density from the remaining context.

The encoder exposes two feature maps:

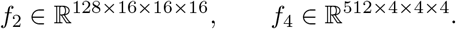

The intermediate feature map *f*_2_ preserves spatial detail at the patch-grid resolution, while *f*_4_ provides compressed global structural context. A lightweight convolutional decoder upsamples the global feature map, fuses it with the spatial feature map, and predicts the voxel values of each masked patch.

The reconstruction loss is computed only over masked patches:

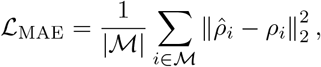

where *M* denotes the set of masked patches, *ρ*_*i*_ is the target patch, and 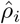 is the reconstructed patch. Each target patch is normalized before reconstruction loss computation so that the loss is not dominated by high-density regions.

We use a curriculum masking strategy to gradually increase reconstruction difficulty. Early epochs use smaller masked blocks and a lower masking ratio, while later epochs use larger contiguous blocks and higher masking ratios. This forces the encoder to move beyond local interpolation and learn longer-range structural context.

To improve robustness to cryo-EM-specific variation, augmentations are applied to the encoder input while the reconstruction target remains clean. These include random 3D flips and rotations, random cropping, simulated missing density, Gaussian blur, local contrast normalization, Gaussian noise, contrast jitter, and sparse density dropout.

After MAE pretraining, Figure 5, only the encoder backbone weights are exported:

**Figure 5.**
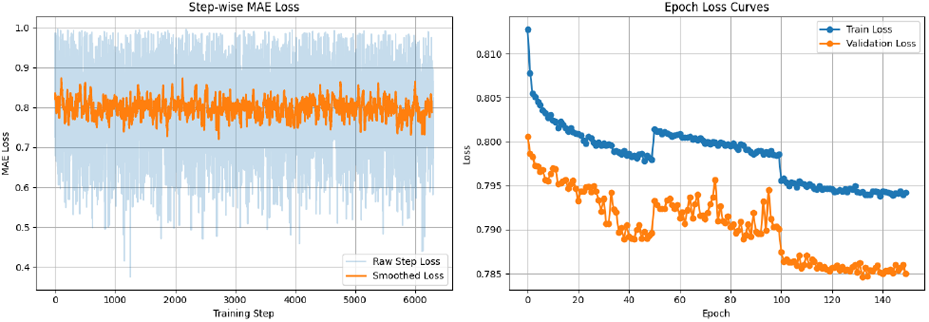
Mae loss curves

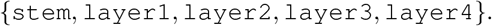

The projection head is not transferred and is instead randomly initialized during cross-modal JEPA training. This ensures that pretraining provides structural initialization for the volume backbone while allowing the shared latent projection space to be learned directly from sequence–structure alignment.

### 3.4. Cross-Modal JEPA Training Objective

After encoding protein sequences and cryo-EM density maps into a shared 512-dimensional latent space, we train the model using a cross-modal JEPA objective. The goal is to learn predictive relationships between modalities while remaining robust to the weak supervision and structural ambiguity inherent in cryo-EM datasets.

Unlike strict contrastive learning methods that assume exact one-to-one correspondence between modalities, the JEPA formulation learns predictive consistency between latent representations. This is particularly important in cryo-EM settings where:

- a single cryo-EM map may contain multiple proteins or subunits,
- the same protein sequence may appear in multiple conformational states,
- and sequence–structure correspondences may be incomplete or noisy.

#### 3.4.1. Bidirectional Cross-Modal Prediction

Given a protein sequence embedding *z*_seq_ and cryo-EM volume embedding *z*_img_, the model learns two predictor networks:

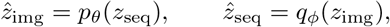

where:

- *p*_*θ*_ predicts the cryo-EM latent representation from the sequence embedding,
- *q*_*ϕ*_ predicts the sequence latent representation from the cryo-EM embedding.

Each predictor consists of a lightweight multi-layer perceptron with LayerNorm, GELU activations, and dropout regularization.

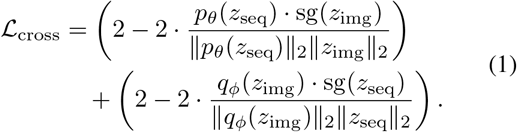

The predictor outputs are L2-normalized before cosine comparison, preventing trivial collapse toward zero-valued embeddings.

This bidirectional prediction objective encourages the latent representations of sequence and structure to contain mutually predictive information without requiring exact embedding equality.

While the JEPA prediction loss trains the predictor networks, it does not necessarily force the raw sequence and image embeddings to be directly aligned in latent space. In practice, we observed that JEPA prediction alone produced weak retrieval performance because predictor networks could compensate for embedding mismatch.

To directly optimize retrieval quality, we introduce a symmetric InfoNCE loss over normalized sequence and image embeddings:

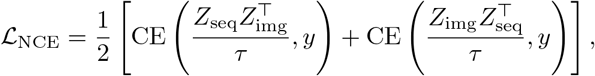

where:

- *τ* is the temperature parameter,
- *y* denotes the diagonal positive-pair labels,
- and cosine similarity is computed between L2-normalized embeddings.

This objective directly optimizes ranking behavior for sequence-to-image and image-to-sequence retrieval, aligning training with the Recall@*k* metrics used during evaluation.

To prevent representational collapse and encourage embedding diversity, we incorporate VICReg-style variance and covariance regularization.

Importantly, these regularization losses are applied to the *raw projection outputs before L2 normalization*. Applying variance regularization after normalization is ineffective because unit-normalized 512-dimensional vectors have extremely small per-dimension variance:

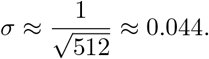

As a result, enforcing a target variance of *γ* = 1 after normalization becomes physically unattainable and causes the variance loss to remain saturated throughout training.

Instead, we compute VICReg statistics on the unnormalized projection outputs:

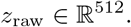

The variance regularization term encourages each embedding dimension to maintain sufficient batch-level variation:

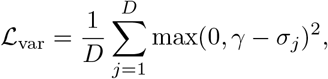

where *σ*_*j*_ denotes the standard deviation of embedding dimension *j* across the batch.

The covariance regularization term penalizes redundancy between embedding dimensions:

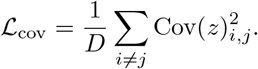

Both losses are computed independently for the sequence and image branches and then summed.

To improve covariance estimation stability under gradient accumulation, raw embeddings are accumulated across multiple micro-batches before VICReg statistics are computed. This substantially improves covariance estimation quality compared to computing VICReg losses on very small per-step batch sizes.

The complete training objective is:

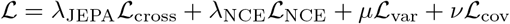

In our implementation, the default weights are:

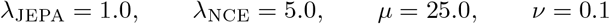

#### 3.4.2. Two-Stage Cross-Species Training

Training is performed in two stages.

The model is first trained on the *Mus musculus* cryo-EM sequence–structure dataset, Figure [6]. The cryo-EM volume encoder is initialized from the MAE-pretrained Cryo2Struct backbone, while the JEPA predictors and sequence projection heads are randomly initialized.

**Figure 6.**
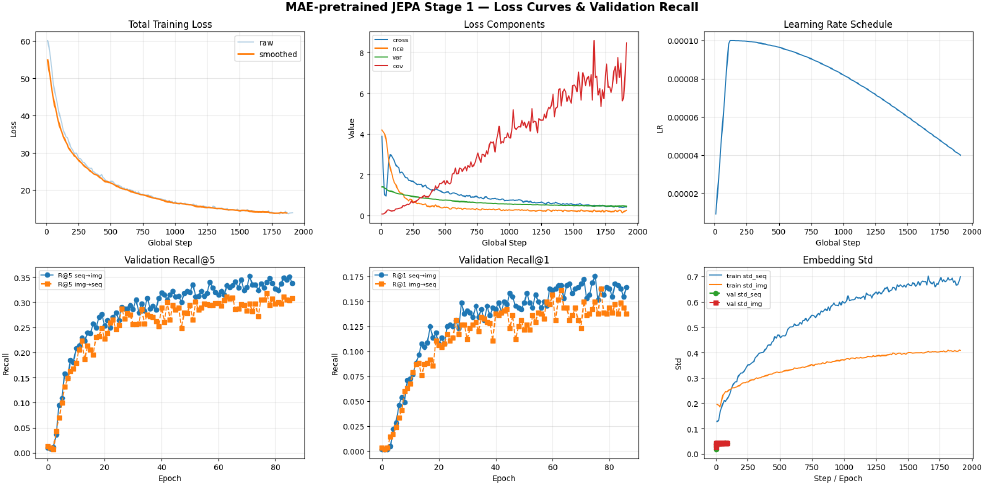
Stage 1: Mouse Training Loss Curves

During this stage, the ESM-2 backbone remains frozen for the first 15 epochs before the top transformer layers are gradually unfrozen using a smaller learning rate.

The best checkpoint from the mouse training stage is then used to initialize the human training stage, Figure [7]. This transfers:

- the cryo-EM volume encoder,
- sequence projection heads,
- partially adapted ESM-2 layers,
- and JEPA predictor networks.

**Figure 7.**
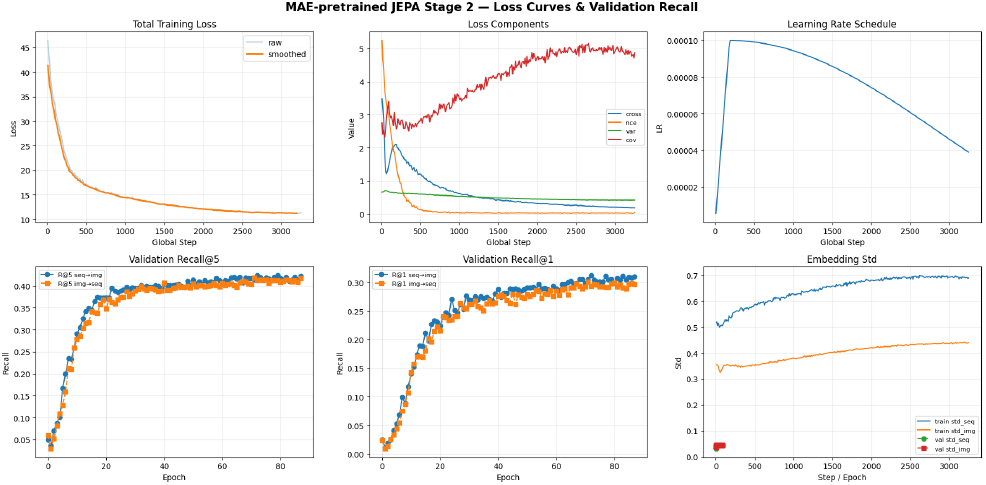
Stage 2:Human Training Loss Curves

Optimizer state and learning-rate schedules are reset for the human dataset to allow adaptation to the new distribution while preserving learned structural priors.

Because the sequence encoder is already partially adapted after Stage 1, ESM-2 unfreezing occurs earlier during human training.

Evaluation is performed using bidirectional retrieval metrics:

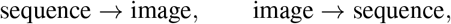

measured using Recall@1 and Recall@5 over the validation set.

## 4. Discussion

Our experiments, Table 1, suggest that the primary bottleneck in multimodal cryo-EM representation learning is not the protein sequence encoder, but rather the ability of the volume encoder to learn meaningful structural representations from sparse and noisy cryo-EM density maps.

**Table 1.** Comparison of retrieval performance across different approaches.

| Approach | R@1 | R@5 | R@10 | MRR |
| --- | --- | --- | --- | --- |
| CLIP ViT Encoder (Baseline) | 0.109 | 0.1813 | 0.2523 | NA |
| CLIP Resnet Encoder (Baseline) | 0.1254 | 0.2255 | NA | NA |
| JEPA baseline (Cosine + EMDB maps only) | 0.003 | 0.016 | 0.032 | 0.019 |
| Multi-loss (JEPA baseline + Combination of losses) | 0.009 | 0.026 | 0.051 | 0.029 |
| Scaled training (Multi-Loss + AlphaFold Augmentation) | 0.030 | 0.120 | 0.181 | 0.083 |
| BYOL without pretraining | 0.1242 | 0.2008 | 0.2688 | 0.2393 |
| BYOL with CT pretraining | 0.2593 | 0.3720 | 0.4511 | 0.4187 |
| BYOL with CT w/ Transfer learning | 0.2613 | 0.3920 | 0.4643 | 0.4332 |
| Pretrained ViT Cryo2Struct MAE w/ Transfer Learning | 0.1423 | 0.2750 | 0.3462 | 0.3035 |
| Pretrained ResNET Cryo2Struct MAE w/ Transfer Learning w/ CT Weights | 0.2887 | 0.4141 | 0.4951 | 0.4432 |
| **Pretrained ResNET Cryo2Struct MAE w/ Transfer Learning** | **0.3024** | **0.4236** | **0.5184** | **0.468** |

The ESM-2 sequence encoder already provides strong biological priors learned from large-scale protein corpora. In contrast, the cryo-EM encoder must learn structural geometry directly from single-channel volumetric densities with comparatively limited supervision. Across nearly all experiments, improvements to the volume encoder produced the largest gains in retrieval performance.

Initial CLIP-style baselines using both ViT and ResNet encoders demonstrated that direct cross-modal alignment alone was insufficient for robust retrieval. While the models learned some degree of sequence–structure correspondence, the learned representations remained relatively weak and unstable.

Introducing BYOL-based self-supervised pretraining substantially improved performance. This indicates that the encoder benefits from first learning intrinsic cryo-EM structural priors before multimodal alignment. Rather than immediately optimizing sequence–structure correspondence, the model first learns local density continuity, volumetric organization, and reconstruction-specific structural patterns directly from cryo-EM data.

These observations reinforce an important intuition throughout our experiments: successful multimodal alignment depends heavily on the quality of the underlying cryo-EM structural representation.

We additionally explored initializing the volume encoder using pretrained MedicalNet weights derived from CT and MRI datasets. The motivation was that generic volumetric priors such as spatial continuity, hierarchical 3D feature extraction, and volumetric geometric understanding may transfer to cryo-EM representation learning.

MedicalNet initialization produced a substantial improvement over randomly initialized volume encoders, suggesting that pretrained volumetric representations provide a strong starting point even across different imaging domains. This observation further reinforced the idea that the volume encoder was the primary bottleneck in multimodal cryo-EM learning. To better model cryo-EM structure directly, we introduced masked autoencoder (MAE) pretraining on Cryo2Struct cryo-EM volumes. This consistently produced the strongest overall retrieval performance and highlighted the importance of cryo-EM-specific self-supervised learning.

Interestingly, MAE pretraining from random initialization slightly outperformed MAE initialized from MedicalNet weights. While initially counterintuitive, this behavior is consistent with the substantial domain gap between cryo-EM density maps and conventional medical imaging modalities.

MedicalNet representations are optimized for CT and MRI statistics, which contain smoother anatomical gradients and very different intensity distributions compared to sparse molecular cryo-EM densities. As a result, MedicalNet initialization introduces priors that the encoder must partially adapt away from during cryo-EM training. In contrast, random initialization allows the entire pretraining process to focus directly on modeling cryo-EM geometry.

Overall, these experiments suggest that cryo-EM representation learning benefits more from domain-specific structural pretraining than from generic volumetric medical imaging priors.

Cross-species transfer learning from mouse to human datasets consistently improved performance across experiments. Since many proteins and structural motifs are evolutionarily conserved between *Mus musculus* and *Homo sapiens*, the model is able to transfer structural priors learned from mouse cryo-EM data into the human retrieval setting. This suggests that cross-species multimodal transfer learning may provide a practical mechanism for overcoming the relatively limited size of experimentally paired cryo-EM datasets.

One notable observation was that the ResNet-based encoder consistently outperformed the ViT-based encoder despite the theoretical advantages of transformer architectures.

This behavior is likely driven by the scale of the data set and inductive bias. The ViT backbone is substantially more data-hungry and relies on large-scale training corpora to learn stable spatial representations. In contrast, the convolutional inductive bias of the ResNet architecture acts as a strong regularizer for cryo-EM data:

- local convolutions naturally capture secondary-structure-scale density patterns,
- translation equivariance improves robustness to spatial variation,
- hierarchical downsampling provides stable multi-scale structural features.

At the current dataset scale, these inductive biases appear more beneficial than the additional expressivity of global self-attention.

## 5. Results

We evaluate the learned multimodal representation space using bidirectional retrieval between protein sequences and cryo-EM density maps. Given a sequence query, the model retrieves candidate cryo-EM maps, and given a cryo-EM map query, it retrieves candidate protein sequences. Performance is reported using Recall@1, Recall@5, Recall@10, mean reciprocal rank (MRR), median rank, and mean rank. Table 2 summarizes retrieval performance on the full human dataset. The model achieves strong bidirectional retrieval, this indicates that the learned latent space captures meaningful sequence–structure correspondence rather than relying on random similarity.

**Table 2.** Retrieval metrics comparison across different evaluation configurations.

| Config | Direction | R@1 | R@5 | R@10 | MRR | MedianRank | MeanRank |
| --- | --- | --- | --- | --- | --- | --- | --- |
| **Config 1: Validation Queries vs Validation Database (n=947) — Generalisation** |  |  |  |  |  |  |  |
|  | Seq → Map | 47.7% | 60.8% | 64.8% | 54.1% | 2.0 | 81.4 |
|  | Map → Seq | 53.7% | 64.8% | 69.4% | 59.2% | 1.0 | 75.8 |
| **Config 2: Validation Queries vs Full Database (n=3275) — Deployment** |  |  |  |  |  |  |  |
|  | Seq → Map | 27.3% | 51.4% | 57.2% | 38.8% | 4.0 | 278.9 |
|  | Map → Seq | 19.7% | 57.6% | 61.4% | 36.8% | 3.0 | 260.1 |
| **Config 3: Full Dataset (n=3275) — Comparison (includes training maps)** |  |  |  |  |  |  |  |
|  | Seq → Map | 36.2% | 58.9% | 63.8% | 46.8% | 3.0 | 226.5 |
|  | Map → Seq | 34.7% | 65.0% | 68.4% | 48.0% | 2.0 | 204.6 |

The rank distribution in Figure 8 further supports this result. Correct matches are concentrated near the top ranks, with a sharp decay after the first few positions. This shows that the model is not only retrieving correct pairs occasionally, but is systematically ranking true matches highly.

**Figure 8.**
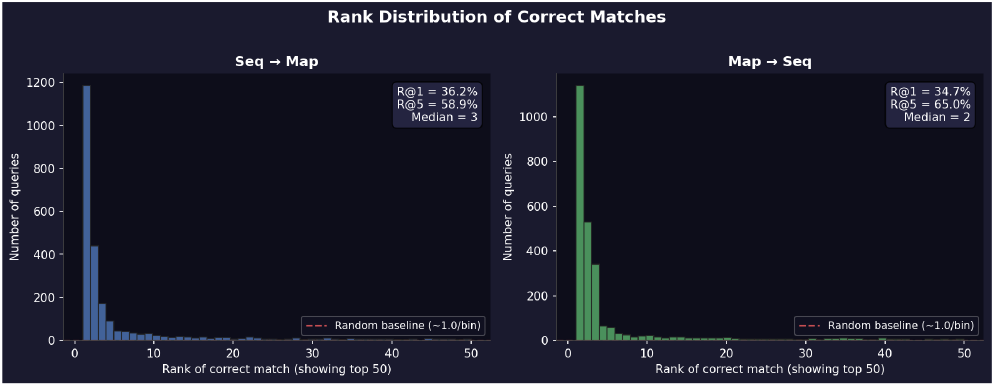
Rank distribution of correct sequence–map and map– sequence matches.

To test whether the learned representation space aligns matched sequence–map pairs, we compare cosine similarities for matched and unmatched pairs. Figure 9 shows a clear separation between the two distributions, with a large alignment gap between true pairs and randomly mismatched pairs. This provides direct evidence that the model learns a shared latent space where paired protein sequences and cryo-EM maps are closer than unrelated examples.

**Figure 9.**
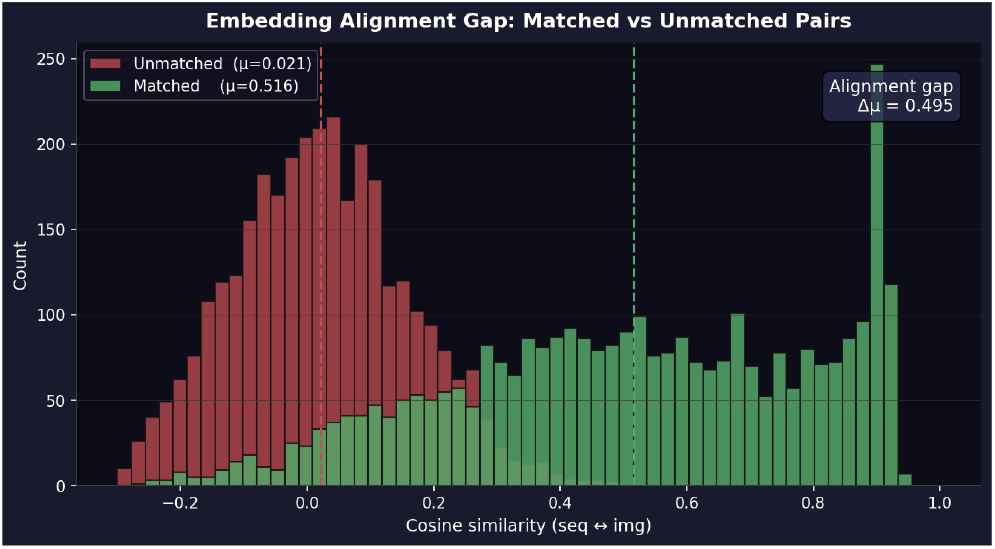
Cosine similarity distributions for matched and un-matched sequence–map pairs.

The similarity heatmap in Figure 10 provides a complementary view of this alignment. The strong diagonal structure indicates that predicted map embeddings from sequence queries are most similar to their corresponding true map embeddings.

**Figure 10.**
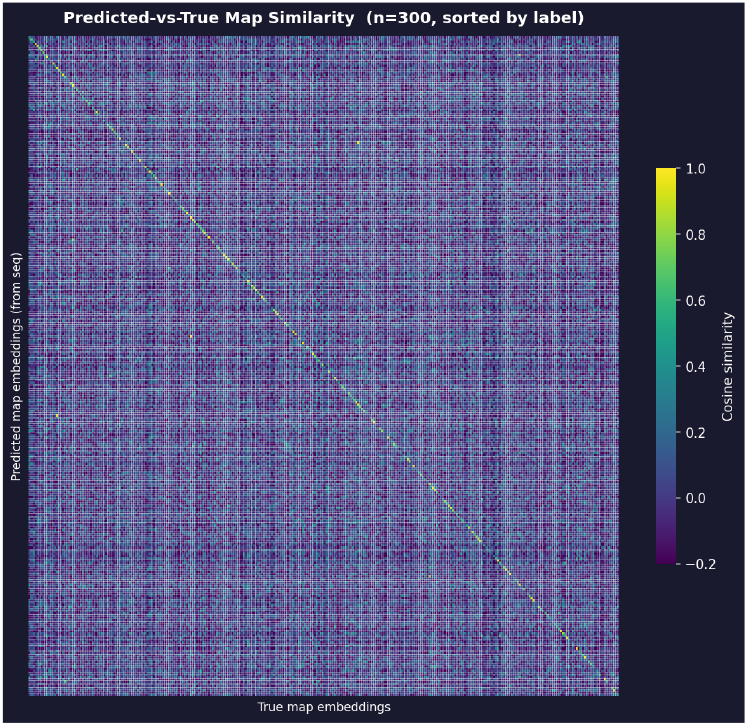
Predicted-versus-true map embedding similarity matrix.

Beyond exact pair retrieval, we evaluate whether the model retrieves maps from the same protein family. Figure 12 shows that top-*k* retrieved maps are strongly enriched for the same protein family compared to a random baseline. This suggests that the learned space captures higher-level biological and structural similarity, not only exact instance-level correspondences.

**Figure 11.**
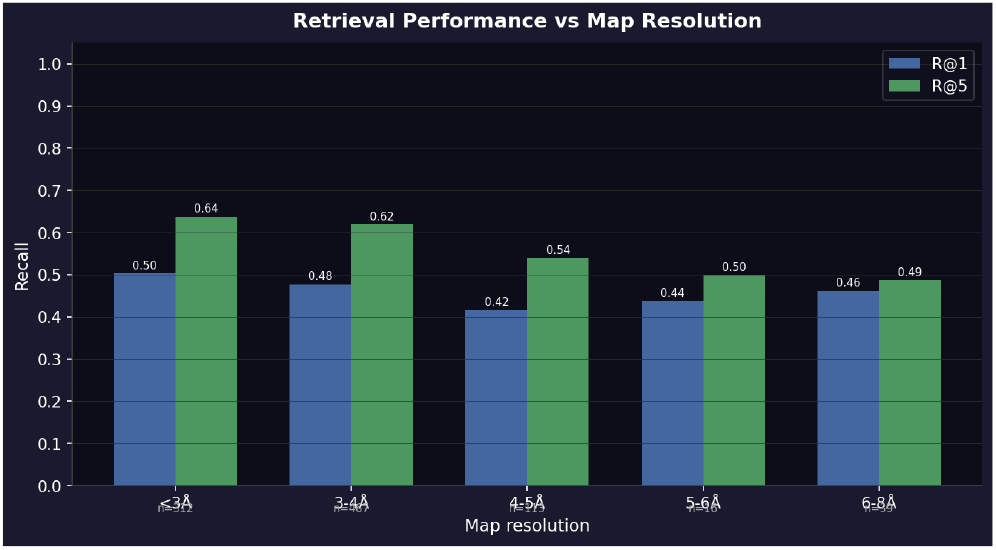
Sequence-to-map retrieval performance binned by cryo-EM map resolution.

**Figure 12.**
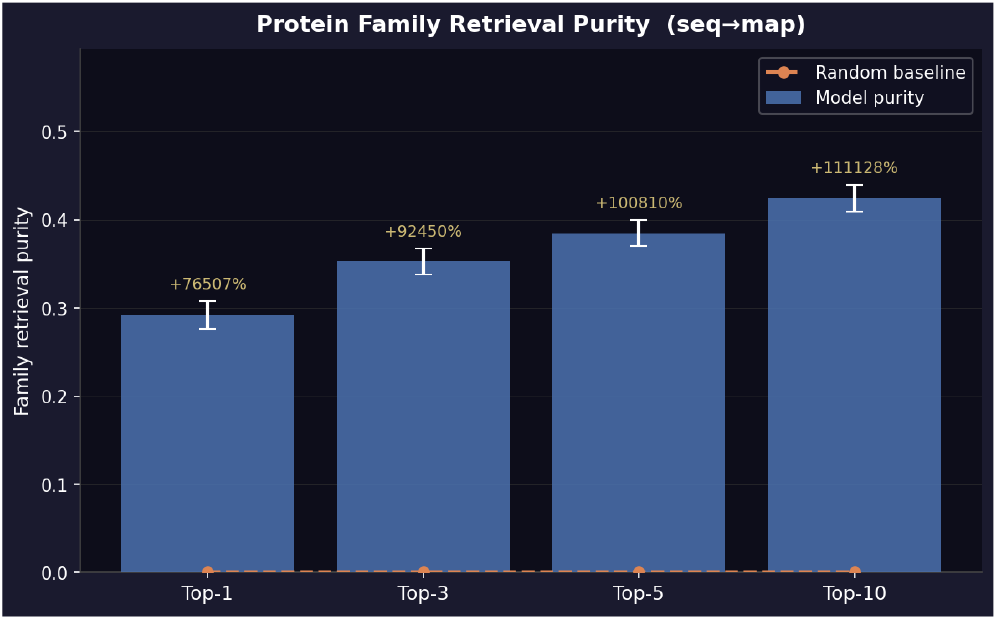
Protein family retrieval purity for sequence-to-map retrieval.

To evaluate transfer beyond the human dataset, we query mouse maps using human sequence embeddings for conserved protein families. Figure 13 shows that the model ranks same-family mouse maps far above random expectation. This supports the hypothesis that the learned latent space captures conserved structural signals that transfer across related species.

**Figure 13.**
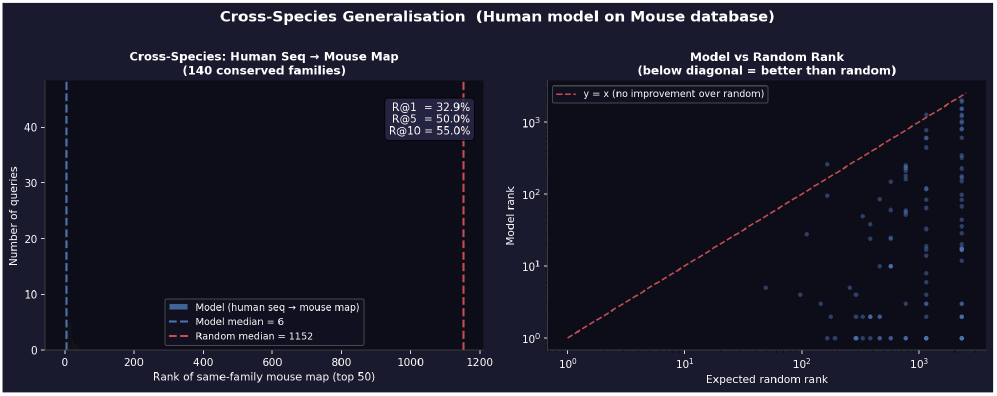
Cross-species retrieval using human sequence queries against the mouse map database.

Figures 14 and 15 show qualitative examples of top-5 retrievals in both directions. In many cases, the correct map or sequence appears among the highest-ranked candidates, and incorrect candidates often remain visually or structurally similar to the query.

**Figure 14.**
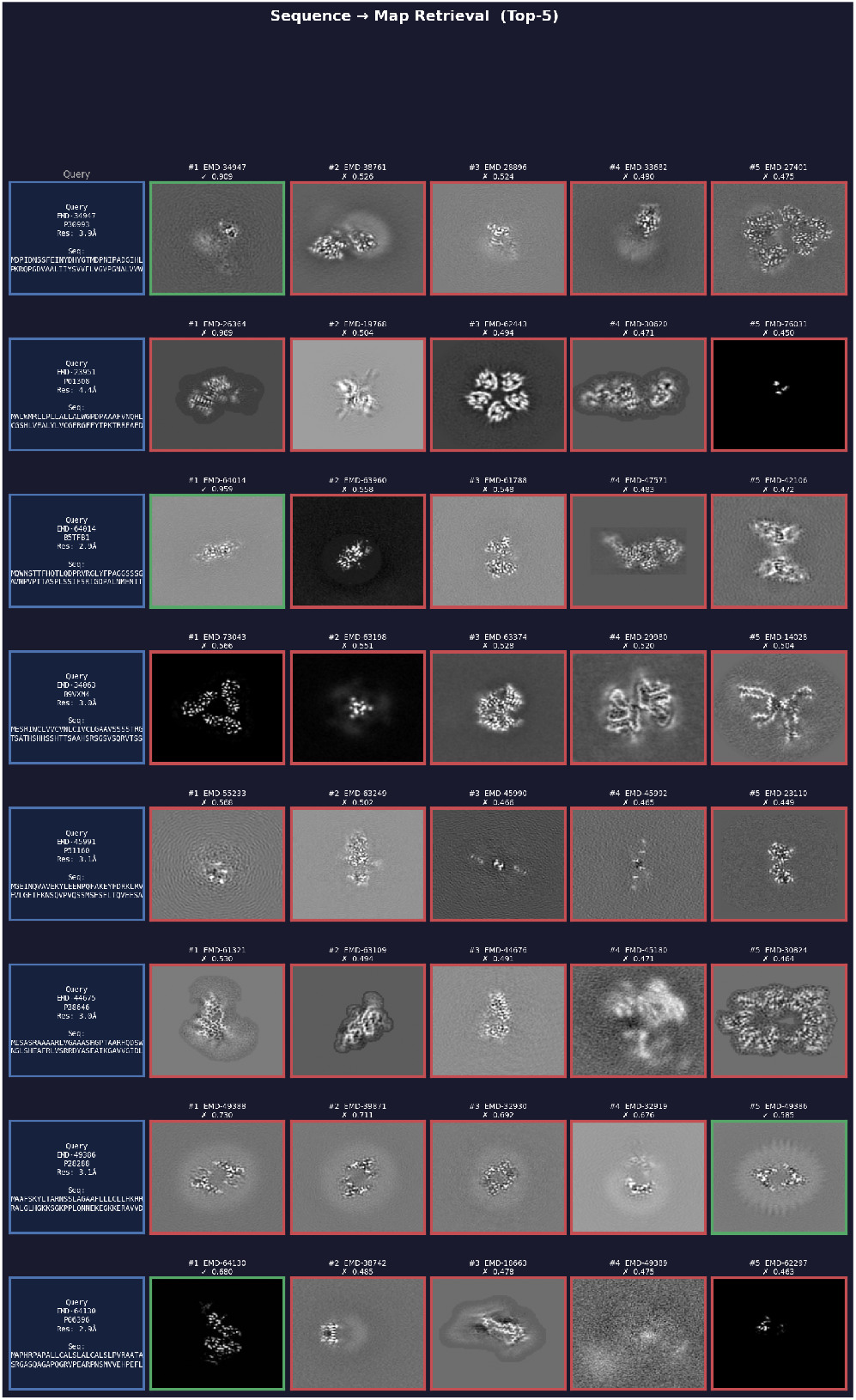
Qualitative sequence-to-map retrieval examples.

**Figure 15.**
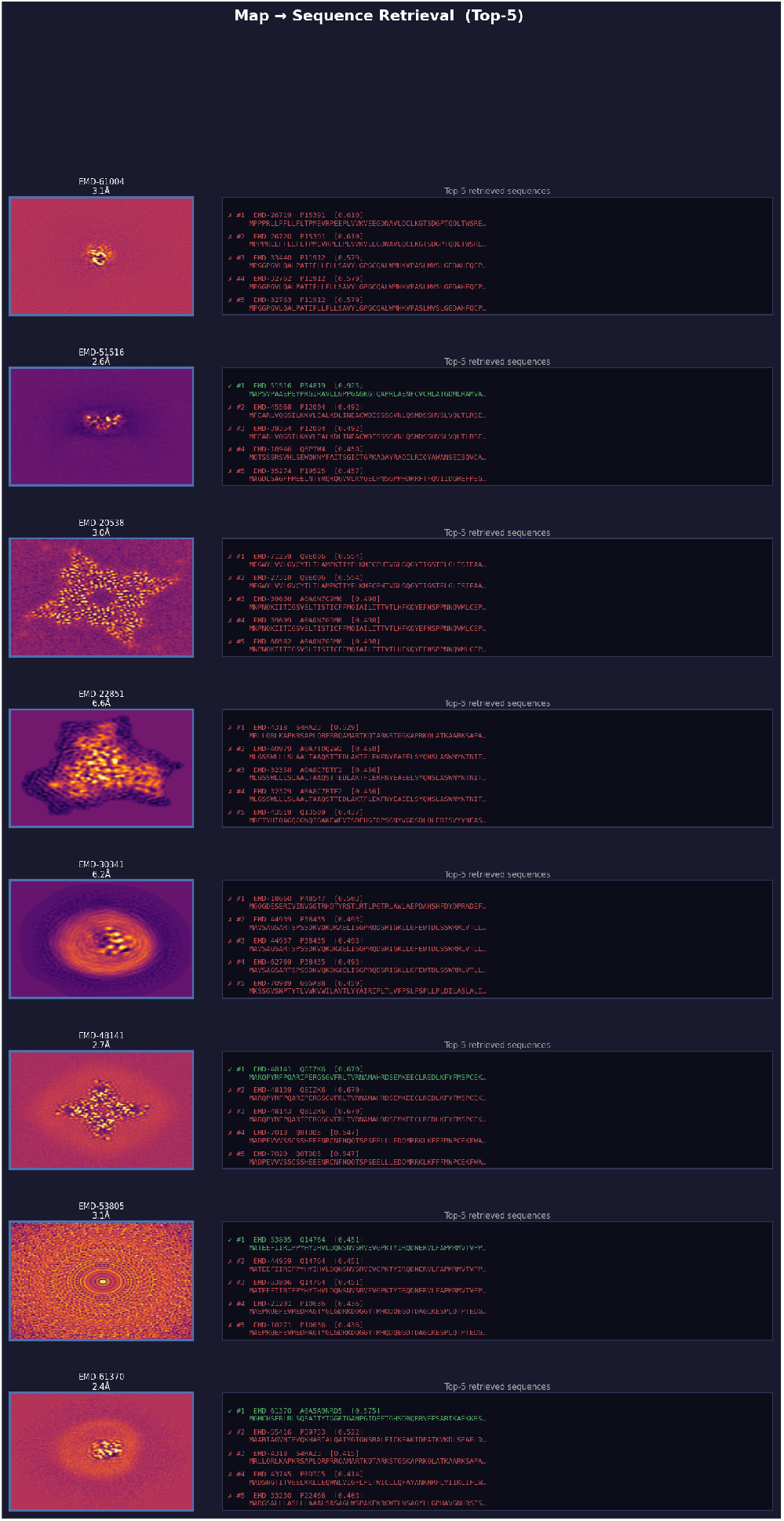
Qualitative map-to-sequence retrieval examples.

Finally, Figure 16 presents the worst sequence-to-map retrieval cases. These failures are informative: the model often retrieves maps with similar visual density structure even when the exact target is ranked poorly. This suggests that remaining errors are not purely random, but reflect limitations of global map-level embeddings when multiple structures have similar density patterns.

**Figure 16.**
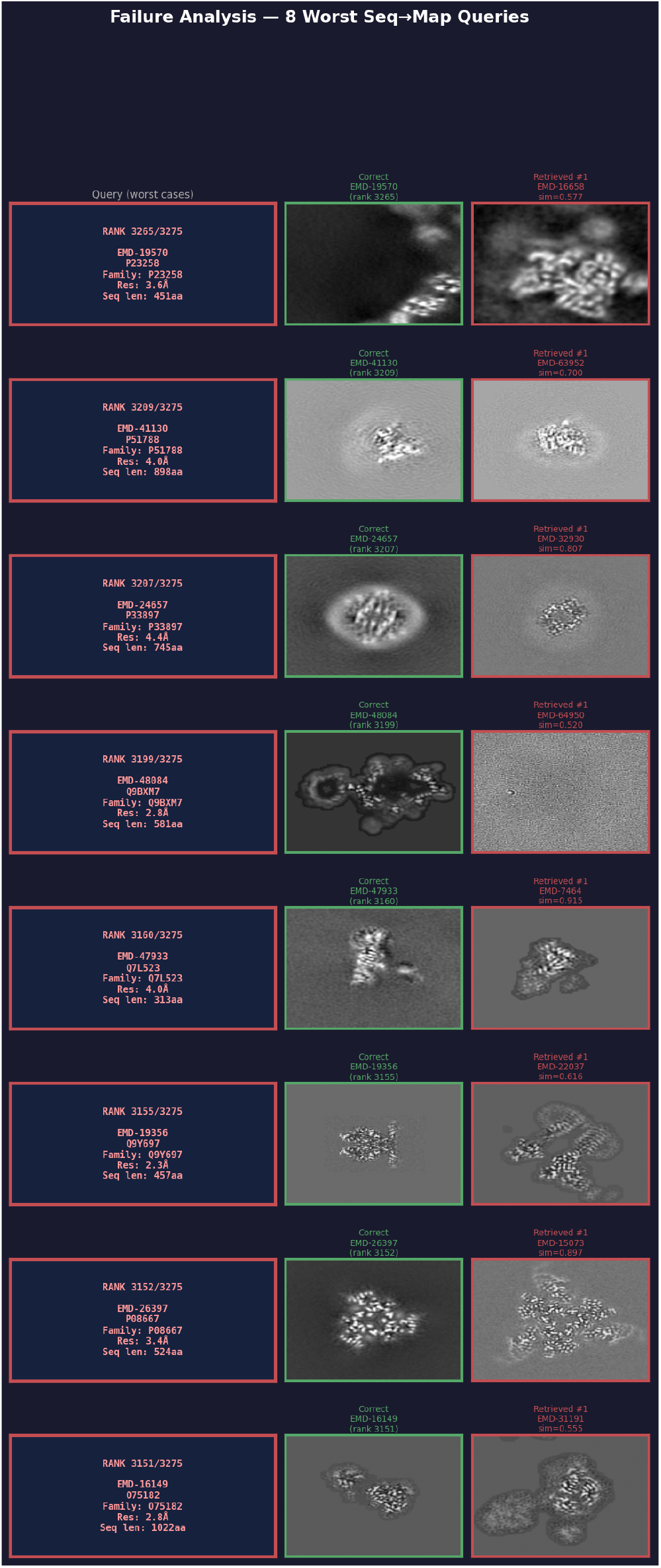
Failure analysis showing worst sequence-to-map retrieval queries.

## 6. Conclusion

Learning shared representations between protein sequences and cryo-EM density maps is an important problem for computational structural biology, particularly in de novo and experimental settings where identifying structures directly from cryo-EM observations remains challenging.

In this work, we presented a cross-modal JEPA-based framework for aligning protein sequences and cryo-EM volumes within a shared latent space. Our experiments show that predictive latent-space learning is more effective than relying purely on strict instance-level matching, especially given the weak supervision, structural heterogeneity, and multi-instance ambiguity present in cryo-EM datasets.

While the current framework demonstrates promising retrieval performance, future work may explore larger-scale cryo-EM pretraining, localized structure-aware representations, and multi-instance modeling for complex cryo-EM assemblies.

Our work suggests that multimodal latent-space learning provides a promising direction toward scalable sequence– structure understanding directly from cryo-EM observations.

